# Multiple functions for the catenin family member plakoglobin in cadherin-dependent adhesion, fibronectin matrix assembly and *Xenopus* gastrulation movements

**DOI:** 10.1101/318774

**Authors:** Glen D. Hirsh, Bette J. Dzamba, Pooja R. Sonavane, David R. Shook, Claire M. Allen, Douglas W. DeSimone

## Abstract

Shaping an embryo requires tissue-scale cell rearrangements known as morphogenetic events. These force-dependent processes require cells to adhere to their neighbors, through cadherin-catenin complexes, and to their extracellular matrix substrates, through integrin-based focal contacts. Integrin receptors are not only important for attachment to the extracellular matrix, but also for its fibrillar assembly. Fibrillogenesis requires actomyosin contractility, regulated in part by cadherin-catenin complexes. One such catenin, plakoglobin, mediates the attachment of actin stress fibers to cadherin cytoplasmic tails through its interactions with actin-binding proteins. In *Xenopus* gastrulae, plakoglobin has been identified as an essential member in the force-induced collective migration of the mesendoderm tissue. In the current study, we have further characterized the role of plakoglobin in two additional morphogenetic processes, epiboly and convergent extension. Plakoglobin-deficient tadpoles are 40% shorter and gastrulae contain notochords that are 60% wider than stage-matched controls, indicating convergent extension defects. The radially intercalating ectoderm of morphant animal caps is nearly twice as thick as controls. Furthermore, morphant embryos exhibit a failure to assemble a fibronectin matrix at the notochord-somite-boundary or along the blastocoel roof. The loss of the fibronectin matrix, while not due to changes in overall patterning, is a result of a failure to assemble the soluble dimers into long fibrils. The force of attachment to a cadherin or fibronectin substrate is reduced in plakoglobin morphants, indicating defects in adhesion to both cadherin and fibronectin. These data suggest that plakoglobin regulates morphogenesis and fibronectin assembly through cell-cell and cell-matrix adhesion.

## Introduction

The extracellular matrix is a common feature of all multicellular organisms and is involved in cell adhesion and signaling, cell migration, cell survival, homeostasis, and maintenance of tissue structural integrity. Spatiotemporal regulation of ECM production and deposition occurs throughout embryonic development resulting in multiple tissues containing different ECM molecules with distinct tensile and viscoelastic properties. These distinct properties ensure proper functioning of the cells, tissues, and organs that interact with assembled ECM. FN is a multifunctional prototypical matrix glycoprotein for which many of the steps involved in fibrillogenesis have been identified. FN is secreted from cells as a dimer, and assembled into fibrils at the cell surface (Rozario and DeSimone, 2010; Schwarzbauer and DeSimone, 2011). Binding of FN to integrin receptors, primarily α_5_β_1_, followed by actin-based contractility applies strain to FN dimers resulting in conformational changes thus exposing cryptic FN-FN binding sites. This contractility and unfolding event is required for fibrillogenesis (Baneyx et al., 2001; Wu et al., 1995; Zhong et al., 1998).

In *Xenopus*, FN is the earliest known matrix glycoprotein expressed in the embryo, with protein translation and fibrillogenesis beginning at the MBT and early gastrulation, respectively. Throughout gastrulation fibrils are assembled onto the blastocoel roof (BCR), serving as a substrate for mesendoderm cell migration (Lee et al., 1984; Winklbauer, 1998) and as a repository of PDGF secreted by BCR cells (Ataliotis et al., 1995; E.Smith et al., 2009). Aside from collective migration of the mesendoderm, FN is also required for radial cell intercalation and epiboly of the BCR tissue (Rozario et al., 2009) and for mediolateral cell intercalation in the dorsal mesoderm during convergent extension (Davidson et al., 2006; Marsden and DeSimone, 2003). FN functions have been a focus of much research, however, most of what is known about steps involved in the expression, secretion, and assembly of FN and other ECM proteins are derived from studies using cell culture monolayers. It is unclear how this translates to assembly along a free surface of a tissue *in vivo*, such as at the BCR.

Cultured cells in a monolayer are relatively flat, maximizing the surface in contact with the substrate. This flattening minimizes the area in contact with neighboring cells. During *in vitro* fibrillogenesis, cell culture dishes provide a substrate to which cells and forming FN fibrils are anchored, and cells rely on this substrate attachment for the generation of forces required for fibrillogenesis (Pankov et al., 2000). During *de novo* FN assembly in a tissue such as along the BCR, cells have no substrate on which to apply forces other than the surfaces of neighboring cells. Calcium-dependent transmembrane adhesion proteins called cadherins mediate most cell-cell adhesions. Cadherin adhesions have been implicated in ECM fibrillogenesis, through kinase cascades as well as force-dependent mechanisms of crosstalk (Dzamba et al., 2009; Weber et al., 2011).

Cadherin-based cell-cell adhesions are divided into two main categories, AJs and desmosomes. In AJs, the actin cytoskeleton is anchored to the cytoplasmic tails of classical cadherins such as E-, N-, and C-cadherin. At desmosomes, keratin or vimentin intermediate filaments (IFs) are anchored to the cytoplasmic tails of desmosomal cadherins, including desmoglein and desmocolin (Saito et al., 2012). C-cadherin (also called EP-cadherin), is the earliest cadherin expressed in *Xenopus* embryos (Ginsberg et al., 1991; Levi et al., 1991). While it is a *bona fide* classical cadherin based on sequence homology and localization at AJs, C-cadherin is also capable of assuming the function of a desmosomal cadherin, interacting with the keratin IF network in mesendodermal cells (Weber et al., 2012). Localization of classical cadherins at desmosome-like structures has been documented in other cell-types including the lens fiber cells of the eye, however, these structures have not been thoroughly explored (Leonard et al., 2008).

The cytoplasmic tails of desmosomal cadherins distinguish their functions from those of classical cadherins, through interactions with different cadherin-binding proteins. The catenin family of proteins is responsible for anchoring cytoskeletal networks at adherens junctions (AJs) and desmosomes (Shapiro and Weis, 2009). β-catenin and plakoglobin (PG; γ-catenin) exhibit high amino acid sequence homology and have overlapping functions. In AJs, β-catenin and PG bind directly to the cytoplasmic tail domains of classical cadherins as well as to α-catenin. α-catenin anchors actin stress fibers at AJs through direct interactions with filamentous actin as well as by recruiting actin binding proteins such as vinculin and α-actinin (Knudsen et al., 1995; Nieset et al., 1997; Peifer et al., 1992). In an analogous manner, PG binds to the tails of desmosomal cadherins and recruits the IF-binding proteins desmoplakin and plakophilin (Garrod and Chidgey, 2008). This function is specific to PG. β-catenin only partially compensates for a loss of PG at desmosomes in PG knockout mice, which eventually die of desmosome-associated heart defects (Bierkamp et al., 1996; Bierkamp et al., 1999; Ruiz et al., 1996).

In addition to their roles in adhesion, β-catenin and PG mediate Wnt signaling. They interact with T-cell factor (TCF) and lymphoid enhancer factor (LEF) to regulate the expression of canonical Wnt target genes (Guger and Gumbiner, 1995; Vonica and Gumbiner, 2007). The LEF-1/PG complex exhibits less nuclear translocation and a lower affinity for DNA compared to the LEF-1/β-catenin complex (Simcha et al., 1998; Zhurinsky et al., 2000), so PG likely plays a more dispensable role in transcriptional regulation. The list of transcripts regulated by β-catenin and PG overlap, with some regulated by both catenins (Martin et al., 2009) and others controlled by either β-catenin or PG alone (Tokonzaba et al., 2013). During *Xenopus* gastrulation canonical Wnt signaling is required for mesoderm induction and axis specification (Fagotto et al., 1997; Guger and Gumbiner, 1995). Overexpression of β-catenin (Wylie et al., 1996) or PG (Karnovsky and Klymkowsky, 1995) on the ventral side of an embryo results in the formation of a second axis, although PG may be acting indirectly (Karnovsky and Klymkowsky, 1995; Klymkowsky et al., 1999; Merriam et al., 1997). Although β- catenin and PG share a high amino acid sequence homology, β-catenin’s main role is in Wnt signaling and tissue patterning. β-catenin morphants exhibit defects in induction of the dorsal organizer, resulting in ventralized embryos that develop no dorsal structures (Heasman et al., 1994). These phenotypes are not seen in PG morphants, whereas β-catenin knockdown has not been reported to disrupt adhesion or the actin cytoskeleton. Another difference is the requirement for PG, but not β-catenin, in desmosomes (Peifer et al., 1992).

Loss-of-function studies in which PG was knocked down in whole embryos have identified the importance of the protein in maintaining cortical actin structures, but further characterization of whole embryo phenotypes were not determined (Kofron et al., 2002; 1997). When PG is knocked down specifically in the collectively migrating mesendoderm tissue, protrusion orientation and persistent migration are perturbed. The requirement of PG in other morphogenetic movements has not yet been reported (Weber et al., 2012).

The major role of PG is at cell-cell adhesions. PG morphant *Xenopus* embryos are flatter in appearance compared to wildtype embryos likely due to a reduction in the formation of actin stress fibers (Kofron et al., 2002). Within the collectively migrating mesendoderm tissue, PG localizes to dynamic desmosome-like structures. In place of desmosomal cadherins desmosome-like structures in the mesendoderm contain C-cadherin, which are attached to keratin IFs through PG. These mechanosensitive protein complexes are essential for mesendoderm cell protrusion polarity and for maintaining the persistent forward migration of the tissue (Weber et al., 2012). The current study further characterizes the role of PG in morphogenesis. Interestingly, PG morphant embryos do not extend along their anteroposterior axis as well as control embryos. This phenotype cannot be explained simply by a defect in collective cell migration and suggests perturbations in other morphogenetic movements. Here we report a role for PG in convergent extension and radial intercalation. The data demonstrate that PG regulates these movements through the assembly of the fibrillar FN matrix. In the absence of PG cell-cell adhesion is perturbed, resulting in defects in matrix assembly.

## Results

### PG regulates convergent extension

Knocking down PG in whole *Xenopus* embryos results in a significant decrease in axial extension (Figure 1 A). PG morphant tadpoles are 40% shorter than controls. Overexpressing a morpholino-insensitive PG RNA construct partially rescues embryo length (Figure 1 A, B). Western blots show that expression of the rescue construct in morphant embryos significantly increases the total level of PG protein

**Figure 1:**
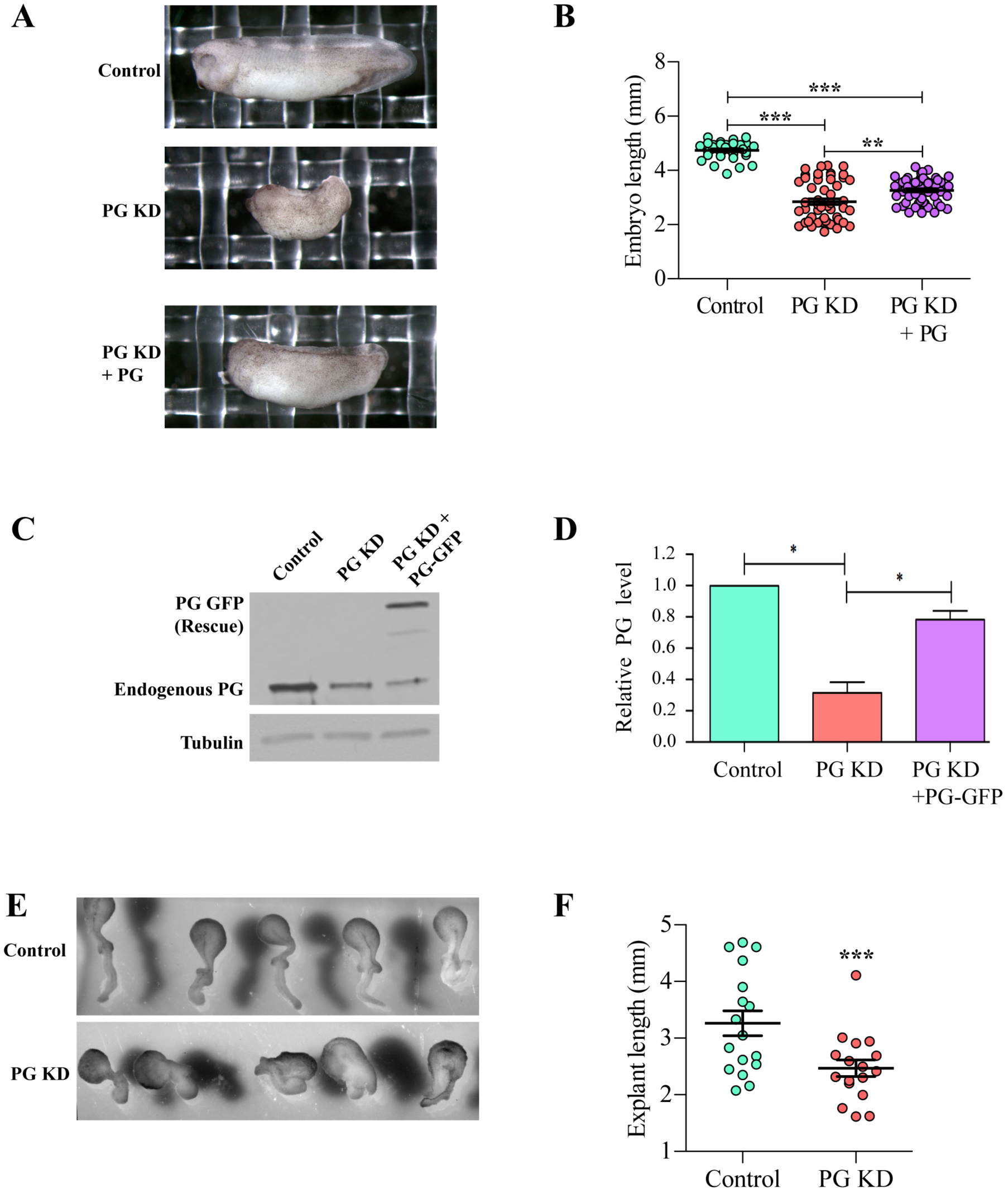
Convergent extension defects in PG morphants. (A) Images representing typical phenotypes of control, PG morphant and PG morphant “rescued” embryos. (B) Quantification of embryo length measurements from control (n=35), PG knockdown embryos (n=53) and PG morphant embryos coinjected with a PG rescue construct that is not targeted by morpholino (n=56). (C) Representative western blot of lysates from whole embryos injected with control morpholino, PG morpholino, or PG morpholino with the nontargetable PG RNA construct (rescue). (D) Quantification of the relative PG expression in control, PG morphant, and “rescue” embryos. Rescue levels are the sum of endogenous PG and the PG-GFP rescue construct (n=4). (E) Keller sandwich explants from control embryos (n=17) and PG morphant embryos (n=17). (F) Quantification of explant lengths from (E). Statistics were calculated based on Wilcoxon matched pairs signed rank test. *p ≤ 0.05, ***p ≤ 0.001. Differences are not significantly different where not specified.

The shortened embryo phenotype seen in Figure 1 A cannot be explained by mesendoderm migration defects alone and suggests perturbations in additional morphogenetic events, namely radial and mediolateral intercalation, that contribute to anteroposterior elongation. During epiboly and convergent extension, cells intercalate between one another, resulting in a thinning of the animal cap and narrowing of the dorsal mesoderm respectively. These processes require the rearrangement of cell-cell and cell-ECM contacts and actomyosin contractility (Keller et al., 2008; Pfister et al., 2016; Skoglund et al., 2008). Keller sandwich explants were used to assess the effects of PG knockdown on convergent extension movements. Keller sandwiches were constructed by dissecting the dorsal sides of two early gastrula stage embryos and sandwiching the inner surfaces together (Keller et al., 1985). Explants elongate as the dorsal tissues converge mediolaterally and extend along the anteroposterior axis. PG morphant explants extend 25% less than their control counterparts (Figure 1 E). This reduction in axial extension in whole embryos and explants is significant (Figure 1 F), and the data are consistent with defects in convergent extension.

During convergent extension, the dorsal mesoderm cells elongate and extend protrusions in the mediolateral direction. Cells use these protrusions to attach to neighbors through cadherin-based adhesions to pull themselves between one another driving intercalation (Keller et al., 2000; Pfister et al., 2016; Skoglund et al., 2008). The axial mesoderm cells that give rise to the notochord and the paraxial mesoderm cells surrounding the notochord increase the length of their mediolateral axis and shorten the width of their anteroposterior axis as they intercalate. These elongated cells in control embryos exhibit high length-to-width ratios of nearly 3:1. In PG morphants, the length-to-width ratios of both axial and paraxial mesoderm cells are significantly reduced to about 1.5:1 (Figure 2 A, B). Morphant notochords are also significantly wider than control notochords, another result of defects in axial extension (Figure 2 A, C). Surrounding the notochord, along the notochord-somite-boundary, is a sheath of ECM containing FN, fibrillin, and laminin. This ECM is necessary for proper mediolateral elongation and intercalation of the dorsal mesoderm cells (Davidson et al., 2004; Parsons et al., 2002; Rongish et al., 1998; Skoglund et al., 2006; Skoglund and Keller, 2007). This notochord sheath is disrupted in PG morphant embryos, exhibiting a reduction in FN and fibrillin staining (Figure 2 D).

**Figure 2:**
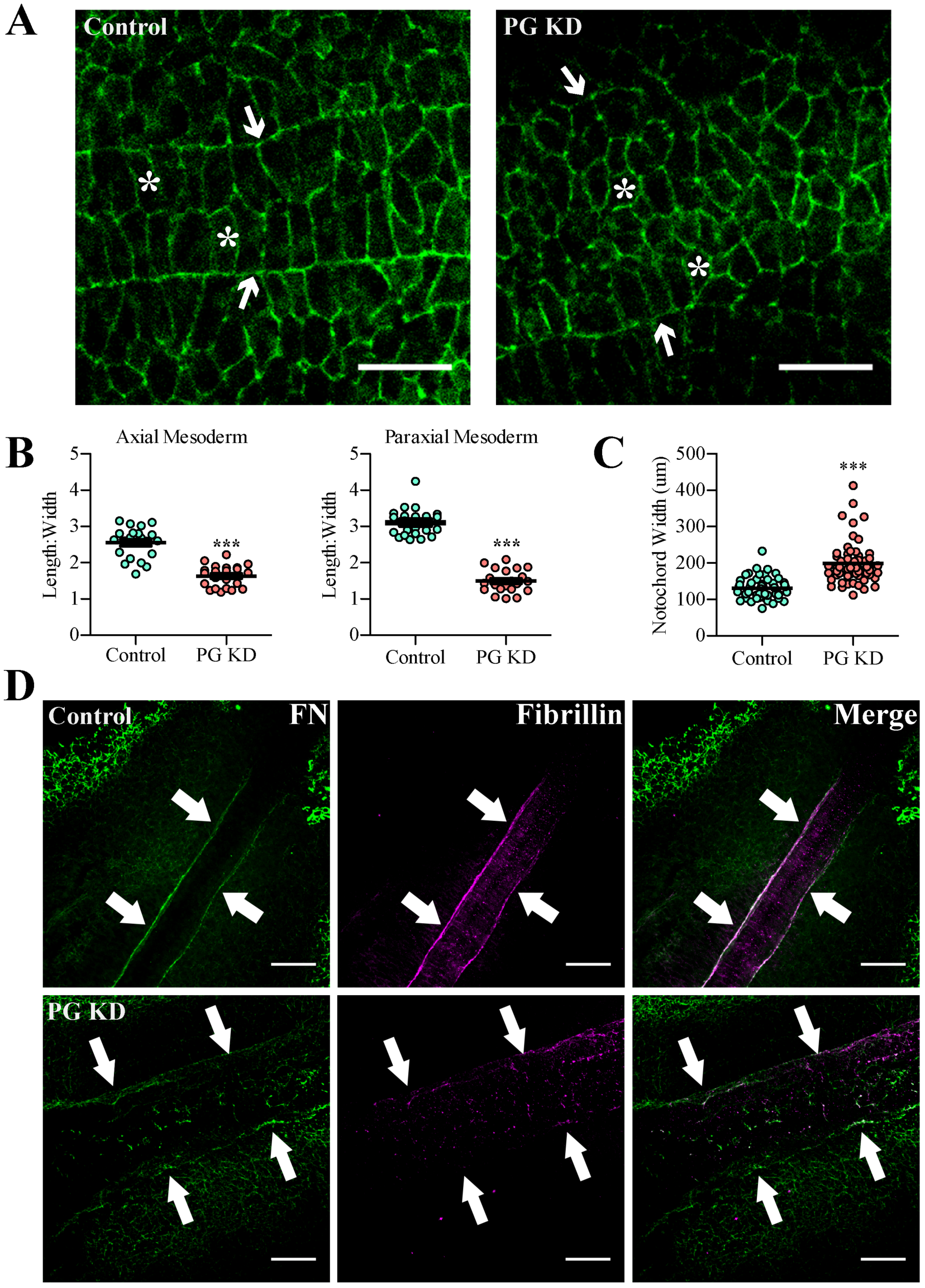
Notochord defects in PG morphants. Representative single confocal sections from control and PG morphant stage 13 dorsal isolates. (A) β-catenin staining (green) outlines cell boundaries. Axial mesoderm cells (asterisks) are shown within the notochord-somite boundaries (arrows) while paraxial mesoderm cells are outside the boundaries. (B) Cell length:width measurements from Axial (n=22 control; n=23 PG KD) and paraxial mesoderm (n=23 control; n=22 PG KD).
(C) Notochord width measurements comparing control and PG morphant dorsal isolates (n=53 control; n=58 PG KD). (D) Dorsal isolates immunostained with antibodies to FN (green) and fibrillin (magenta). Arrows indicate the notochord-somite boundary. Statistics were calculated based on Mann-Whitney tests. ***p ≤ 0.001. Scale bars are 50 μm in (A) and 100 μm in (B).

### PG is a regulator of radial intercalation

When maternal PG is knocked down, morphant embryos exhibit a delay in blastopore closure, suggestive of defects in epiboly (Kofron et al., 1997). This intercalation-driven event thins and spreads the tissue and is required for blastopore closure. During gastrulation, the BCR thins as the deep ectoderm cells intercalate radially between one another and orient cell divisions in the horizontal plane (Marsden and DeSimone, 2001; Rozario et al., 2009). Much like the converging and extending mesoderm, radial intercalation during epiboly requires directed cell migration (Damm and Winklbauer, 2011; Szabó et al., 2016) and cadherin-mediated cell-cell adhesion (Babb and Marrs, 2004; Shimizu et al., 2005). To determine if PG is required for radial intercalation we bisected control and PG morphant embryos and measured the thickness of the BCR tissue. By mid-gastrula stage the BCRs of control embryos have thinned to just two cell layers, measuring about 35 μm thick. However, BCRs of PG morphants remain more than twice as thick and multilayered, with rounded cells protruding into the blastocoel cavity (Figure 3 A, B). This rough appearance along with a reduction in basolateral cadherin accumulation in morphant embryos is suggestive of cell-cell adhesion defects, which may contribute to epiboly defects.

**Figure 3:**
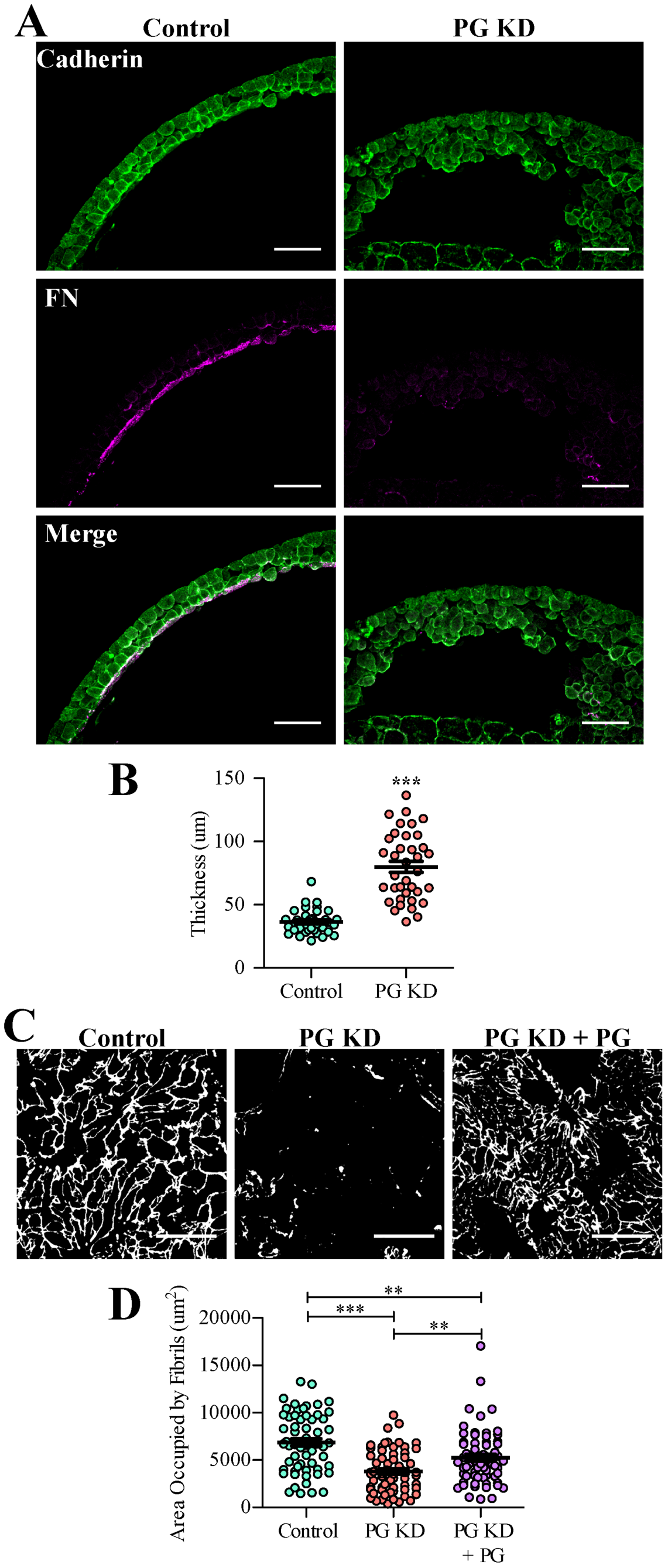
Radial intercalation is disrupted in PG-deficient embryos. (A) Representative transverse sections of stage 11 blastocoel roofs from control (left) and PG KD (right) embryos. Fixed and bisected embryos immunostained for C-cadherin (green), and FN (magenta). (B) Quantification of blastocoel roof thickness from control (n=37) and PG morphant embryos (n=37) (C) Stage 11 animal caps from embryos injected with control or PG morpholino, or with PG morpholino and a morpholino-insensitive PG rescue construct and immunostained using antibodies to FN (white). (D) The area occupied by fibrils on animals caps as in (C) was quantified (n= 61 control, n=64 PG KD; n=63 PG KD + PG embryos). Statistics were calculated based on Wilcoxon matched pairs signed rank tests. ***p ≤ 0.001. Scale bars in (A) are 100 μm and (C) are 15 μm.

In addition to cell-cell adhesion defects, PG-deficient embryos exhibit a reduction in FN assembly along the BCR (Figure 3 A). This is consistent with the reduced FN and fibrillin staining at the notochord-somite-boundary (Figure 2 B). For high magnification imaging with single-fibril resolution, immunostained animal caps were dissected and visualized *en face*. The BCRs of control cells are coated with a dense network of FN fibrils. This network of fibrils is drastically reduced in PG morphants. This fibril loss is rescued by the overexpression of a morpholino-insensitive PG construct (Figure 3 C, D).

### Localization and expression patterns of PG during development

To confirm that PG protein was present at the right time and place to explain the disruption in epiboly and convergent extension in PG morphants, and to compare its expression to that of β-catenin, we used western blots and whole mount immunofluorescence microscopy. Western blots of whole embryo lysates indicate that PG is expressed at relatively low levels pre-MBT (stage 6) and begins to accumulate gradually at MBT (stage 8) and through gastrulation (stages 10-13; Figure 4 A). In contrast, β-catenin is already present at moderate levels by MBT, but expression does not increase until later stages of gastrulation (stages 12-13; Figure 4 B). These data are similar to a previous report that focused on time points that span embryonic development (DeMarais and Moon, 1992) instead of discrete stages within gastrulation.

**Figure 4:**
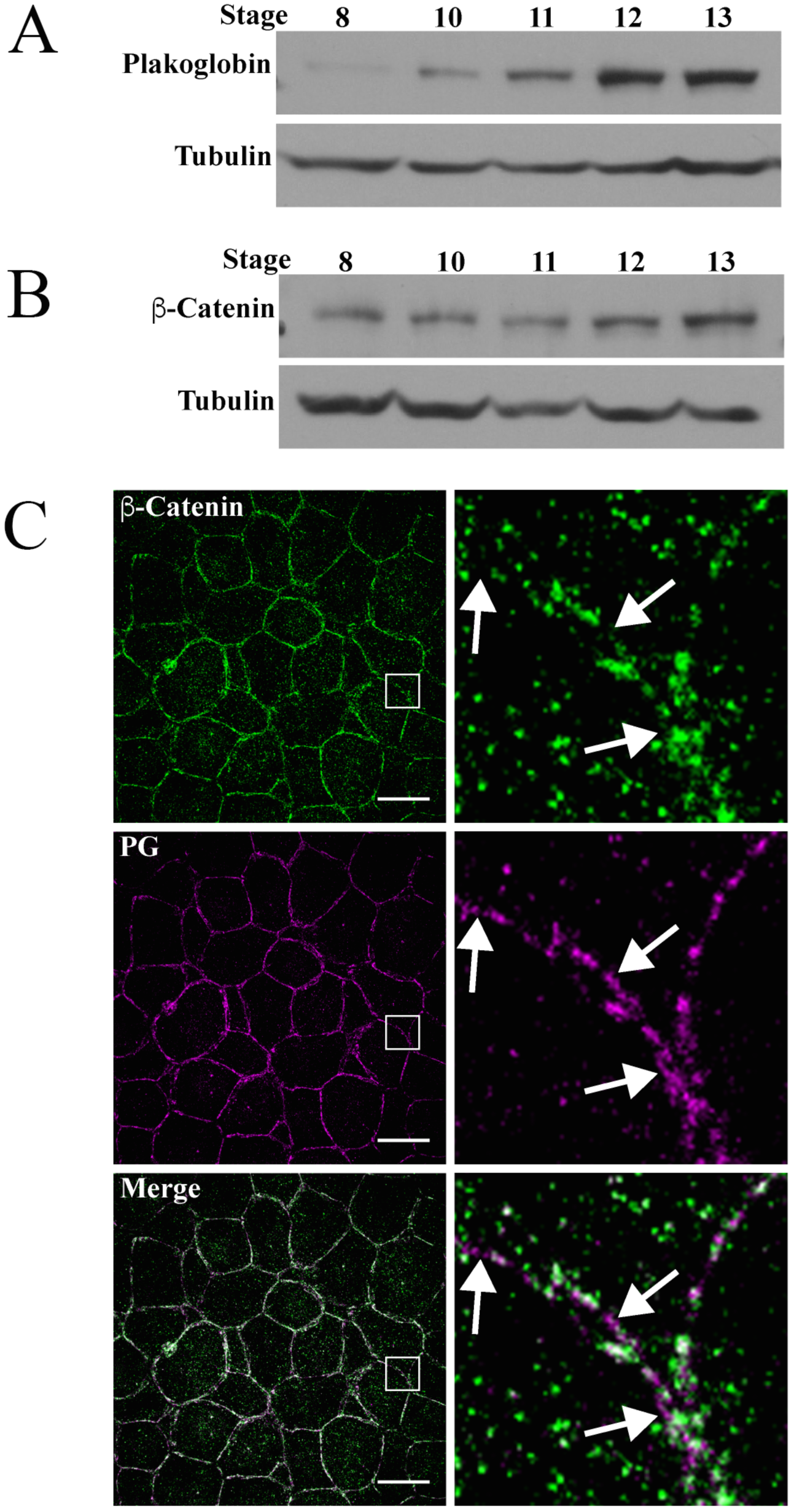
Expression and localization of PG protein during gastrulation. A representative western blot showing expression of PG (A) and β-catenin (B) at MBT (stage 8) and throughout gastrulation (stages 10-13). Tubulin was used as a loading control. (C) Projected confocal Z-stack image of a stage 10 animal cap immunostained with antibodies to β-catenin (green) and PG (magenta). Boxes in the left column images outline the area magnified in the right column. Arrows indicate areas where PG is present without β-catenin. Scale bar is 20 μm.

Spatially, PG and β-catenin colocalize at cell contacts in some cells while remaining distinct in others, such as at the blood-brain barrier (Lampugnani et al., 1995; Liebner et al., 2000). Using confocal microscopy, we found that in early gastrula stage animal cap cells, β-catenin and PG primarily colocalize at punctae along cell boundaries. Separate from these areas of colocalization, PG displays a “beads-on-a-string” appearance adopted by desmosomal proteins as they decorate IFs at nascent desmosomes (Jones and Grelling, 1989; Todorovic et al., 2014) (Figure 4 C). β-catenin and PG primarily localize to cell membranes along the BCR, consistent with a major role for the proteins in this tissue is in adhesion. Although PG and β-catenin serve redundant functions at AJs, their spatiotemporal expression patterns differ in *Xenopus* tissues.

### Mesoderm patterning is normal in PG-deficient embryos

Like β-catenin, PG overexpression is sufficient to induce dorsalization and secondary axis formation, phenotypes associated with canonical Wnt signaling (Karnovsky and Klymkowsky, 1995; Klymkowsky et al., 1999). However, unlike βcatenin knockdown, a loss of PG does not result in ventralization (Kofron et al., 1997, 2002). To confirm the absence of patterning defects in PG morphants, *in situ* hybridizations and immunofluorescence staining of embryos were performed. *Xenopus Brachyury (XBra)* (Smith et al., 1991) and *Chordin (chrd)* (Sasai et al., 1994) were expressed in the early mesoderm and chordomesoderm, respectively, in both control and PG-deficient embryos (Figure 5 A, B). Additionally, immunostaining of dorsal isolates using the 12101 antibody to label the presomitic mesoderm (Kintner and Brockes, 1985), revealed that the tissue remained properly patterned in PG morphant embryos (Figure 5 C).

**Figure 5:**
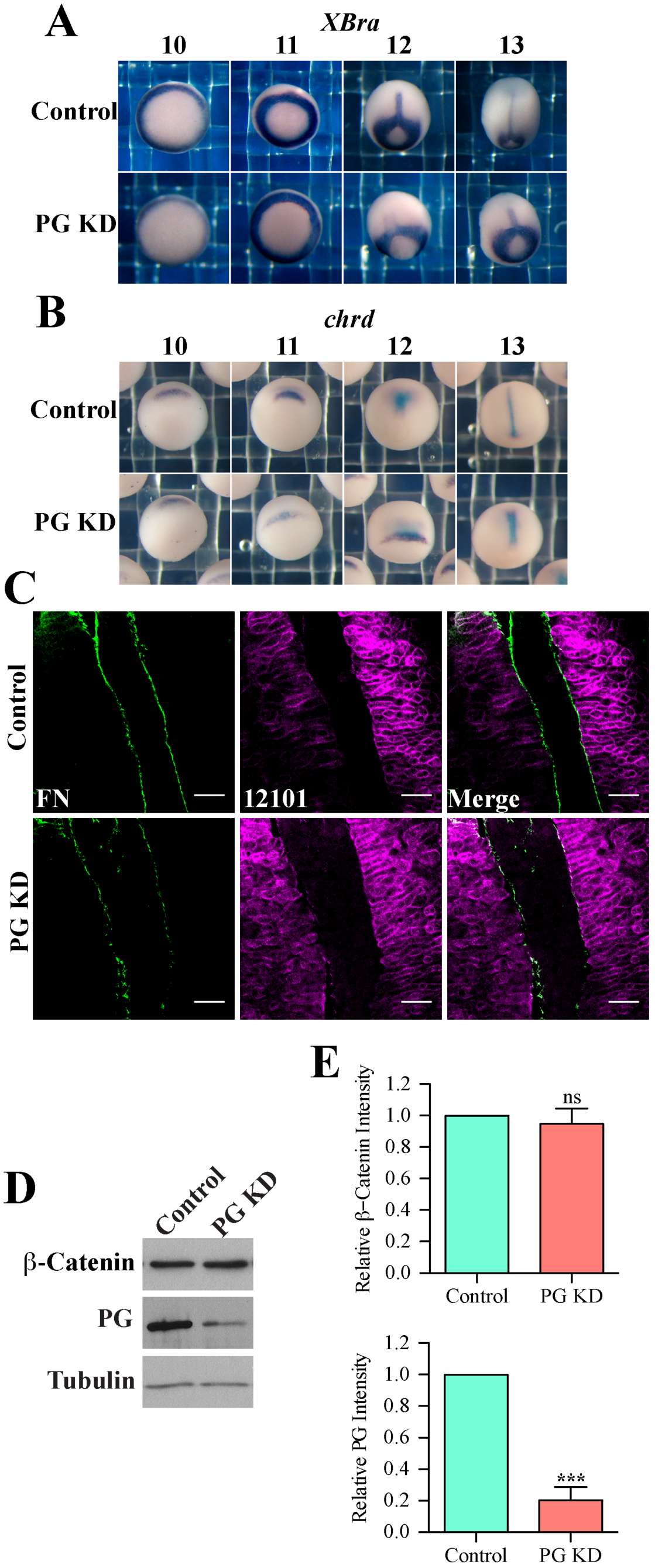
Mesoderm patterning is maintained in PG-deficient embryos. (A, B) *In situ* hybridization of Control and PG morphant embryos showing *Xenopus Brachyury*(A) and *Chordin* (B) expression patterns in during gastrulation. (C) Dorsal isolates stained for FN (green) and presomitic mesoderm marker 12101 (magenta). (D) Representative western blot of whole embryo lysates showing β-catenin and PG, expression. Tubulin was used as a loading control. (E) Comparison of β-catenin and PG protein levels in control embryos (n=13) and embryos treated with PG morpholino (n=13). Statistics were calculated based on Wilcoxon matched pairs signed rank test. ns = not significant, ***p ≤ 0.001. Scale bars in (C) are 50 μm wide.

Because cytoplasmic pools of β-catenin and PG are targeted by the same APC/GSK-3 destruction complex (Rubinfeld et al., 1995), altering the levels of one protein can result in subsequent changes in the other (Li et al., 2007). Densitometric analyses of western blots indicate that β-catenin protein levels are maintained when PG is knocked down (Figure 5 D, E). Overall, data indicate that the observed phenotypes in PG morphants are more likely the result of defects in cell adhesion rather than Wnt signaling.

### FN protein levels are reduced in PG morphant embryos

Although PG knockdown does not affect β-catenin protein levels or cell fate, it does result in reduced FN deposition at the notochord-somite-boundary as well as along the BCR. A loss of fibrils can be caused by a decrease in the FN protein level in the embryo, defects in fibrillogenesis, or a combination of the two. Western blot analysis confirmed a reduction in FN protein levels in PG deficient embryos during gastrulation (Figure 6 A). At the end of gastrulation (stage 13), there is a 40% reduction in total FN protein in PG morphants compared to control embryos (Figure 6 B).

**Figure 6:**
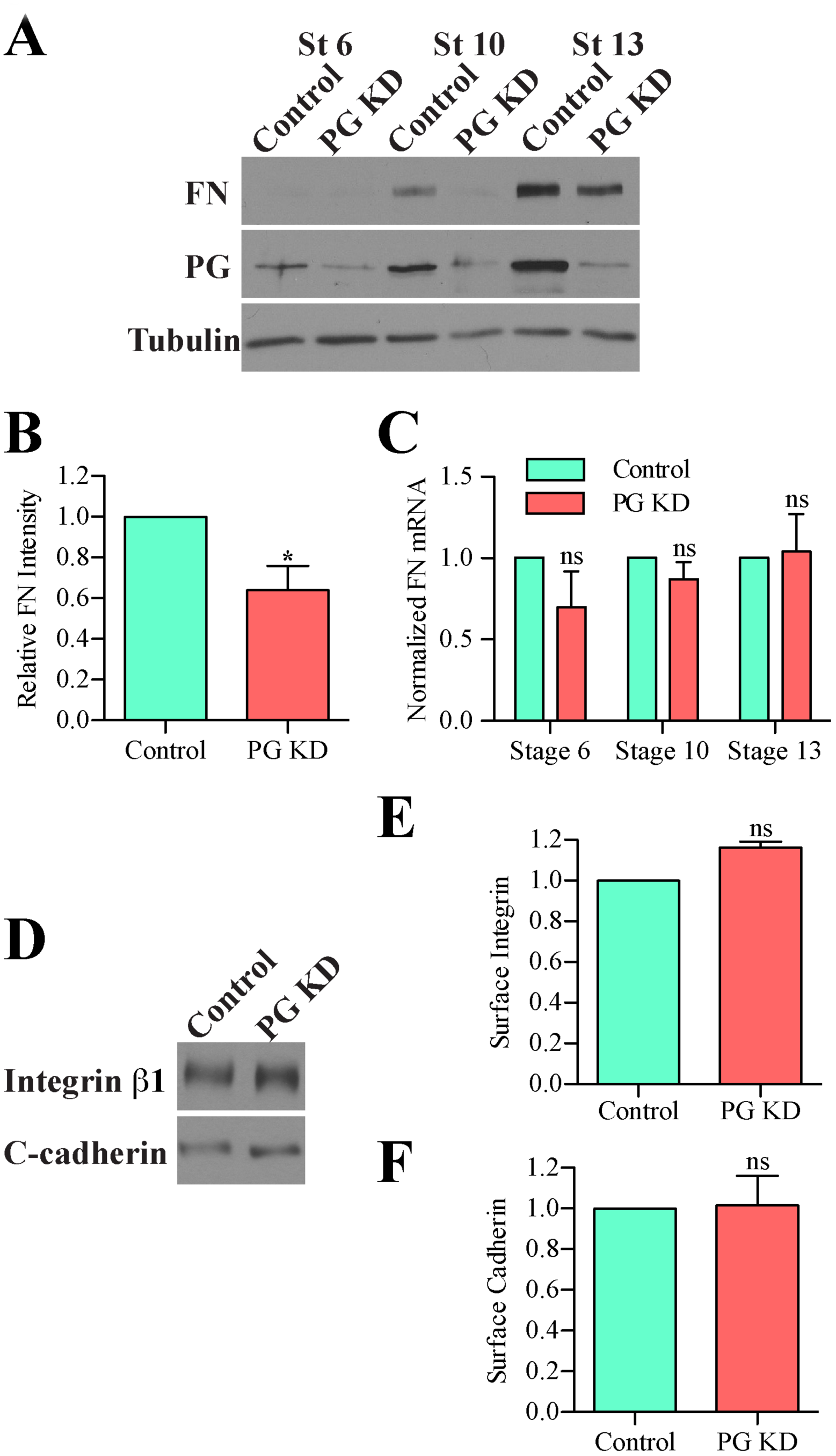
FN protein levels are reduced in PG morphants. (A) Western blots of control and PG morphant whole embryo lysates from stages 6, 10, and 13. (B) FN protein levels relative to tubulin (n=13 control; n=PG KD). (C) Comparison of FN mRNA levels as determined by absolute quantification methods of qPCR (n=3 control; n=3 PG KD). Values within a given experiment were normalized to the level of FN mRNA in control embryos. (D) Surface biotinylated proteins were pulled down with NeutrAvidin agarose and run on an SDS-PAGE gel. Western blots for integrin β1 and cadherin were performed on the samples. (E) Densitometry analyses of integrin β_1_ (E) and cadherin (F) blots in (D) to determine surface integrin and cadherin levels. Statistics were calculated based on Wilcoxon matched pairs signed rank tests on samples from three individual clutches of embryos. ns= not significant. Statistics were calculated based on Wilcoxon matched pairs signed rank test. ns = not significant, **p ≤ 0.01.

One mechanism by which PG may regulate the FN protein is through transcription or transcript stability. Β-catenin has been shown to regulate FN transcription in *Xenopus laevis* through canonical Wnt signaling (Gradl et al., 1999). PG is also reported to function in transcriptional regulation in some cell types (Aktary et al., 2013; Maeda et al., 2003; Tokonzaba et al., 2013). Knocking down PG results in rapid degradation of FN mRNA (Todorović et al., 2010). Quantitative reverse transcription PCR (qPCR) shows that FN transcript levels are maintained in PG morphant embryos, suggesting that PG does not regulate FN transcription or mRNA stability at this stage in development (Figure 6 C). Thus PG morphants have decreased FN translation or protein stability.

### Integrin and cadherin surface protein levels are maintained in PG morphants

Cells in the dorsal mesoderm and the BCR of PG morphants are rounded, suggesting cell adhesion defects. One possible reason for these defects in adhesion as well as the reduced FN assembly along the notochord and BCR is a decrease in cell surface receptors involved in these processes. Cell-cell adhesions in *Xenopus* gastrulae are mediated by C-cadherin, while integrins are responsible for assembling secreted FN dimers into fibrils. Integrin and cadherin receptors are trafficked to the cell surface where they perform their functions in cell adhesion. The catenin family of proteins can physically interact with the uncleaved pro-cadherin in the golgi apparatus and endoplasmic reticulum and are thought to regulate trafficking of the cadherin receptors to the cell membrane (Chen et al., 2009; Curtis et al., 2009; Wahl et al., 2003). Cell-surface biotinylation was used to address the possibility that PG is involved in cadherin or integrin trafficking. Whole animal caps surface proteins were labeled with non-cell-permeable biotin, lysed, and precipitated using NeutrAvidin agarose. Densitometric analyses of western blots indicate equivalent integrin β_1_ and C-cadherin surface levels in control and PG morphants (Figure 6 D-F). This confirms that changes in cell-cell adhesion or FN assembly state are not due to reductions in surface C-cadherin or integrin levels.

### PG is required for FN assembly

Cadherin overexpression is sufficient to induce precocious FN assembly through an actomyosin-based mechanism. This suggests that cell-cell adhesions perform an important role in fibrillogenesis (Dzamba et al., 2009). PG is one of the proteins involved in anchoring the actin cytoskeleton to cadherin adhesions and may regulate fibrillogenesis through the same mechanism. In order to study fibrillogenesis in PG morphants without affecting whole embryo FN protein levels, we knocked down PG in a subset of cells while maintaining expression in the rest of the embryo. PG morpholino was injected into one animal cap cell at 32-cell stage (stage 6) and *de novo* FN fibrillogenesis along the BCR was visualized at stage 11 (mid-gastrulation). Adjacent to the BCR, the blastocoel cavity contains soluble FN dimers (Lee et al., 1984). Therefore, knockdown cells remain exposed to an equivalent FN concentration as their adjacent uninjected counterparts (Figure 7 A). Fixed and immunostained animal caps reveal that FN fibrillogenesis is perturbed in PG morphant cells, while the surrounding uninjected cells are able to assemble FN into fibrils (Figure 7 B). In a similar experimental approach, PG was overexpressed in a subset of animal cap cells by injecting PG mRNA animally into 1 cell at the 4 cell-stage. Embryos were fixed at stage 10 (early gastrulation). At this stage of development, cells do not normally assemble FN into fibrils (Dzamba et al., 2009). Cells overexpressing PG were able to assemble FN fibrils precociously (Figure 7 C), a phenotype similar to that of cadherin overexpression (Dzamba et al., 2009).

**Figure 7:**
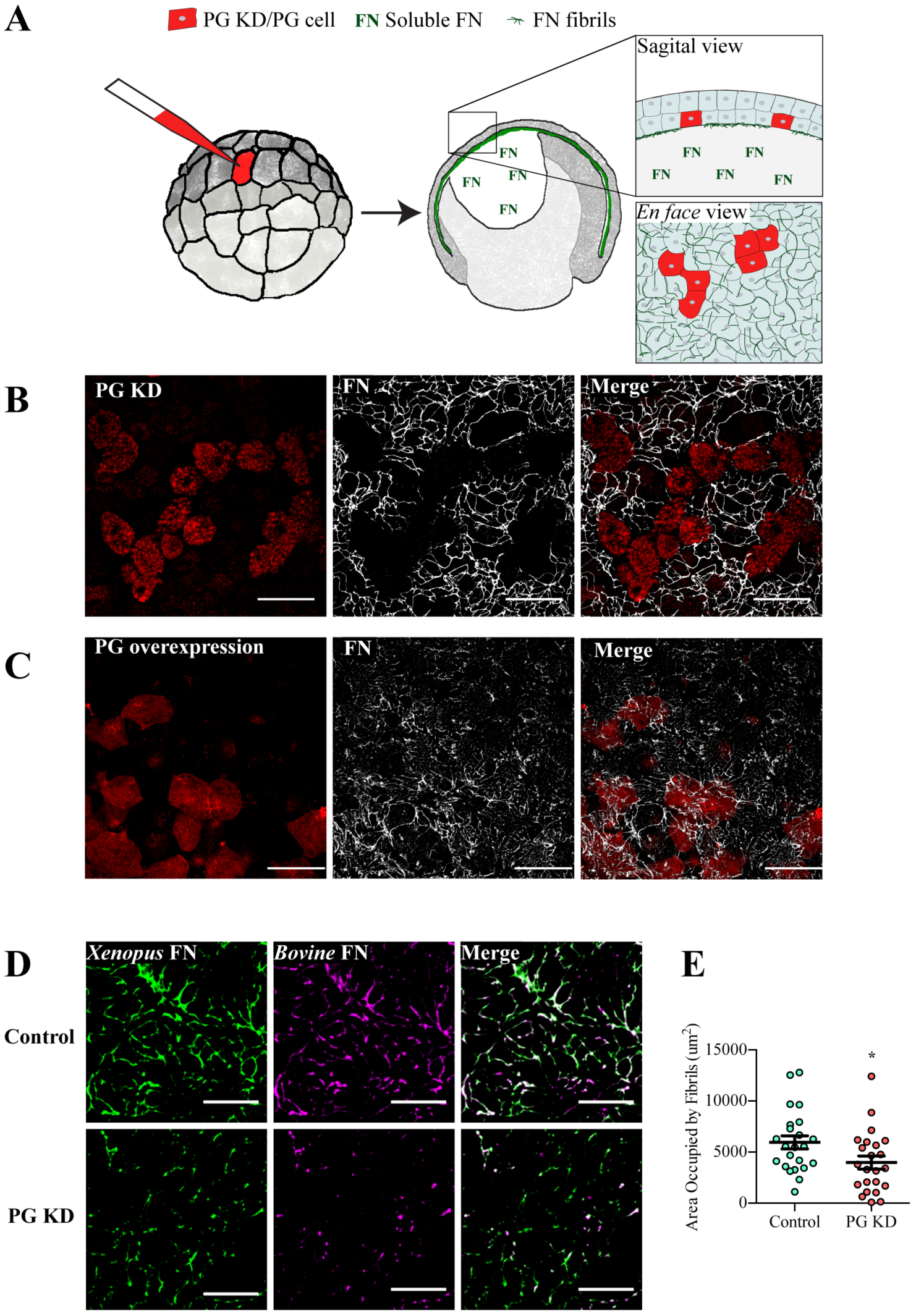
PG is required for FN assembly. (A) Cartoon showing experimental design used in (B). Embryos at 32 cell stage were injected with PG morpholino into a single blastomere (red). At gastrulation, cells within the BCR are exposed to the same pool of soluble FN in the BC. Animal caps were isolated and immunostained for FN fibrils (green and white). (B) Stage 11 animal caps from embryos coinjected with dextran and PG morpholino into 1 of 32 cells. (C) Stage 10 animal caps from embryos co-injected with PG mRNA and fluorescent dextran (red) in 1 of 4 cells. (D) Animal caps from control or PG morphant embryos injected with *Bovine* FN (magenta) and stained for endogenous *Xenopus* FN (green). (E) Area occupied by *Bovine* FN fibrils in (D) was quantified (n=23 control; n=23 PG KD). Statistics were calculated based on Mann-Whitney tests. *p ≤ 0.05, **p ≤ 0.01, ***p ≤ 0.001. Scale bars are 50 μm (B and C) and 15 μm (D)

Knocking down PG in a small subset of animal cap cells leads to a loss of fibrillogenesis in a cell autonomous manner (Figure 7 B). The BCR cells are thought to assemble soluble FN dimers from the blastocoel fluid. One possible explanation for the decrease in assembly is that the local concentration of soluble FN is decreased at the surface of the morphant cells. To avoid this possibility while testing for the requirement of PG in fibril assembly we knocked down PG in whole embryos and reintroduced exogenous FN protein. Embryos were injected with PG morpholino immediately following fertilization. Prior to gastrulation, a fluorescently-tagged *Bovine* FN protein was injected directly into the blastocoel cavity of morphant embryos. At stage 11, animal caps were dissected, fixed, and endogenous FN identified using an antibody that recognizes *Xenopus* but not *Bovine* FN. Control embryos are able to assemble fibrils containing both exogenous *Bovine* FN and endogenous *Xenopus* FN and show extensive overlap between the two species of protein. PG morphant embryos were unable to assemble *Xenopus* or *Bovine* FN into fibrils, confirming a defect in fibrillogenesis (Figure 7 D, E).

### PG morphant cells do not form strong adhesions to cadherin or FN

Crosstalk between cadherin and integrin-based adhesions has been identified in both cell culture and *in vivo* systems (Dzamba et al., 2009; Langhe et al., 2016; Weber et al., 2011). It is possible that through direct mechanical linkage or indirect signaling pathways, the observed defects in fibronectin fibrillogenesis in PG morphants are a result of reductions in adhesion strength. To address the possibility that PG is involved in attachment to FN we performed atomic force microscopy (AFM). A single mesendoderm cell was attached to a Cell-Tak-coated cantilever and brought into contact with cadherin or FN substrates. After a short five-second incubation, cells were detached from the substrates as adhesive strength was measured. Force-distance curves indicate a reduction in the maximum adhesive strength to both cadherin and FN substrates in PG morphant cells when compared to controls (Figure 8 A-C). To test for specificity of decreased attachment strength, forces associated with attachment the second heparin-binding domain of FN (Hep II) (Ramos and DeSimone, 1996) and non-specific attachment to poly-L-lysine (PLL) and were measured. Mesendoderm cells attach to a Hep II substrate via syndecan receptors rather than integrins, therefore, Hep II serves as a receptor-mediated control for attachment. PLL serves as a control for non-receptor-mediated attachment. There is no significant difference in the force of attachment between control and PG morphants on a Hep II substrate (Figure 8 D) or the PLL substrate (Figure 8 E) indicating that the reduction in adhesion to cadherin and FN is integrin dependent (Figure 8 B, C).

**Figure 8:**
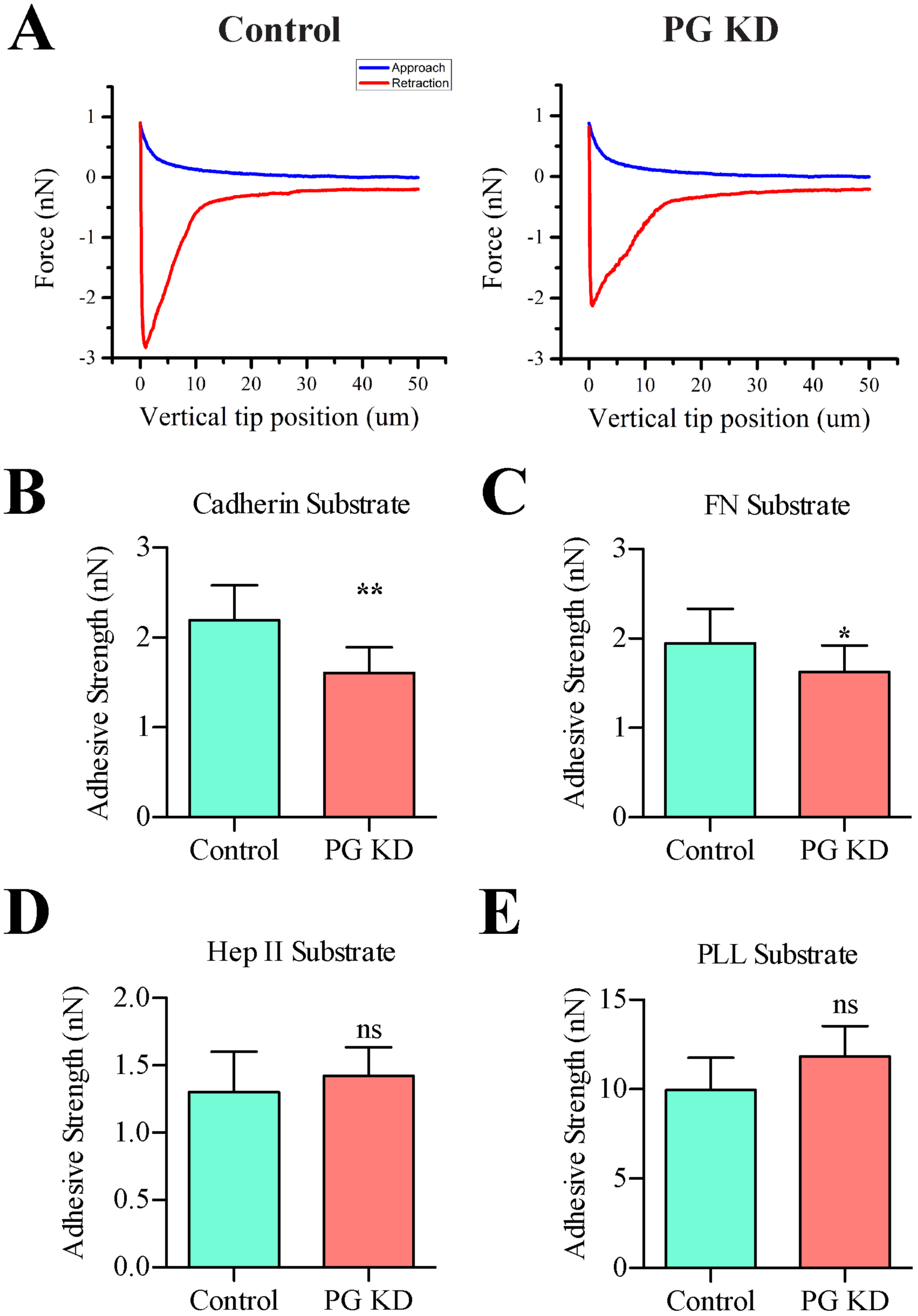
Adhesive strength to cadherin and FN is reduced in PG-deficient cells. (A) Representative force-distance curves of single cell adhesion experiments on a C-cadherin substrate. Single control or PG KD cells were attached to cantilevers and allowed to attach to C-cadherin (B), FN (C), HepII (D), or poly-l-lysine (E) substrates. Approach and retraction curves are shown in blue and red, respectively. Adhesive strength was determined for 10-12 cells per condition for each substrate. Cells from three independent clutches were analyzed for each substrate. Statistics were calculated based on Wilcoxon matched pairs signed rank test. ns = not significant, *p ≤ 0.05, **p ≤ 0.01.

## Discussion

Gastrulation is a complex process that involves numerous coordinated tissue movements. These morphogenetic events rely on ECM assembly and cellular adhesion. The importance of cadherins in cell-cell adhesion and integrins in attachment to and assembly of an ECM has been well documented. However, there is still much that we do not understand about the coordination of these two processes. In this study, we have further characterized the role of PG, a cadherin-associated protein, in the context of gastrulation and morphogenesis. Our data indicate that PG plays an essential role in cell adhesion and FN assembly.

*In vivo* loss-of-function studies on PG-null mice and maternal-RNA KD *Xenopus* embryos reveal that PG anchors actin stress fibers and keratin IFs at AJs and desmosomes, respectively. PG knockout mouse embryos die as a result of heart defects associated with a loss of desmosomal integrity (Ruiz et al., 1996; Bierkamp et al., 1996). When maternal PG mRNA is depleted in *Xenopus* oocytes using antisense oligonucleotides, embryos exhibit a loss of cortical actin stress fibers, mild adhesion defects, and significant developmental delays (Kofron et al., 2002; 1997). In contrast, knocking down PG after fertilization allowed embryos to develop through initial cleavage stages before the protein levels were reduced, and we were able to study the function of PG in embryos with a milder phenotype. We previously identified the recruitment of PG and keratin to C-cadherin in response to force as a regulator of mesendoderm cell polarity and migration (Weber et al., 2012). The current study demonstrates that PG is involved in additional tissue movements during gastrulation including convergent extension and epiboly through its effects on cell-cell and cell-matrix adhesion and FN fibrillogenesis.

In the current study, we have found that PG plays an essential function in morphogenesis through its role in cell adhesion and FN fibrillogenesis. Short morphant embryos exhibit defects in convergent extension and epiboly (Figures 1-3), two processes that require coordination of cell-cell and cell-matrix adhesion (Babb and Marrs, 2004; Keller et al., 2008; Marsden and DeSimone, 2001; Pfister et al., 2016; Rozario et al., 2009; Shimizu et al., 2005; Skoglund et al., 2008). PG morphants have normal surface integrin β_1_ and C-cadherin levels (Figure 6), but do not form strong attachments to cadherin or FN substrates (Figure 8) and do not efficiently assemble FN into fibrils (Figure 7). Reports indicate mechanisms of crosstalk between cadherin and integrin-based adhesions (Bazellières et al., 2015; Marsden and DeSimone, 2003; Mertz et al., 2012) at least in part through Rac and Pak-mediated actomyosin contractility (Dzamba et al., 2009). The contractile force generated at cadherin-based adhesions is transduced through the actin cytoskeleton to integrin adhesions. Force applied to FN dimers induces a conformational change, exposing cryptic FN-FN interacting domains leading to fibril extension (Zhong et al., 1998). Our data suggests that PG is essential for the transmission of the force required for FN assembly.

The function of PG protein in the context of embryonic development has not been well studied. We have identified a role for PG in the assembly of a FN ECM. The function of PG in FN fibrillogenesis is likely through anchoring of the actin cytoskeleton to AJs and the transmission of actomyosin contractility (Figure 9). The cortical actin network is connected to actin filaments at cadherin/catenin AJs as well as to integrins in focal adhesions. As the two halves of a FN dimer are bound to integrins on adjacent cells, actin dependent forces move the integrins centripetally along the basal surface leading to unfolding of FN to exposing cryptic FN-FN binding sites that allow for fibril assembly. In the absence of PG, cadherin-mediated adhesion is perturbed (Figure 8 A, B) the actin cytoskeletal network is disrupted (Kofron et al., 2002) and subsequent attachment strength to FN is reduced (Figure 8 C). Together, our data suggest that PG is indispensable for matrix assembly, a process required for morphogenesis and embryonic development.

**Figure 9:**
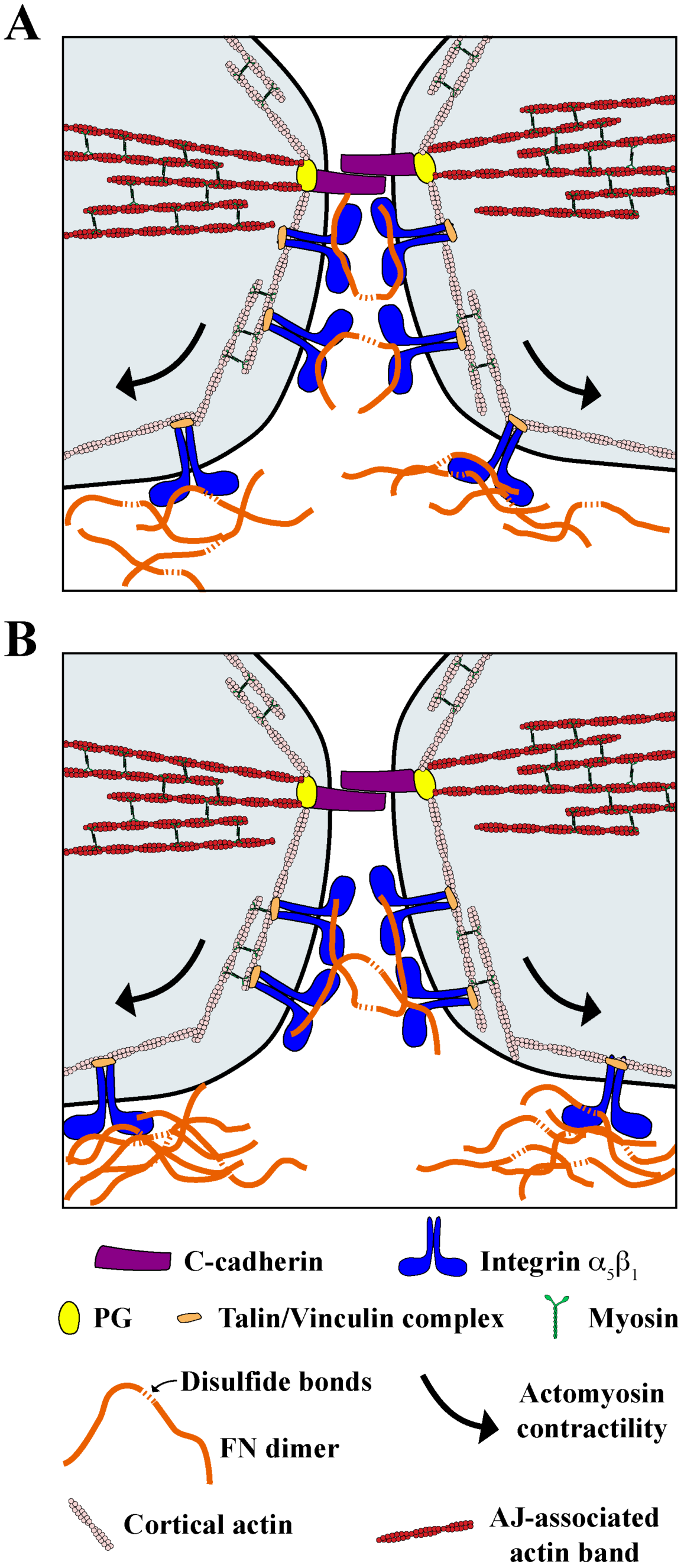
Model of *in vivo* FN assembly. A model depicting FN assembly over time (A-B). Adherens junctions between animal cap cells are mediated by C-cadherin (purple). PG (yellow) recruits actin-binding proteins to anchor actin filaments (red) to the cadherin adhesions. Actin contractility is mediated by myosin (green). α_5_β_1_ integrins (blue) link extracellular fibronectin dimers (orange) to cortical actin stress fibers in the cell. Myosin-based contractility drives the centripetal movement of integrin receptors (arrows), applying strain to the bound FN dimers. Force-unfolded FN dimers bind to one another to form a network of fibrils.

## Materials and Methods

### Embryos and explants

*Xenopus laevis* embryos were obtained and fertilized *in vitro* using standard methods and staged according to Nieuwkoop & Faber (1967). Keller sandwiches were prepared as described in Keller et al. (1985). Single cell experiments were performed using mesendoderm and animal cap cells that were dissociated in calcium- and magnesium-free Modified Barth’s Saline (MBS) (Sive et al., 2000).

### Morpholino and RNA constructs

Fertilized embryos were injected with morpholinos or RNA constructs into both hemispheres of stage 2 embryos or into 1 blastomere located on the animal cap at the 4 or 32 cell stage as indicated. Morpholinos were injected into whole embryos at a concentration of 25 pg/embryo or into 1 blastomere at 32 cell stage at a concentration of 0.78 pg/embryo. RNA was injected into whole embryos at a final concentration of 500 pg/embryo and into 1 blastomere at 32 cell stage at 15.6 pg/embryo. Injections were performed in a solution of 3% Ficoll in 0.1x MBS and incubated in 0.1x MBS until the specified developmental stages (Sive et al., 2000). Standard control (CCTCTTACCTCAGTTACAATTTATA) and PG morpholinos (TTTCCACTACGTCTCCCAAATCCAT) (Weber et al., 2012) were purchased from Gene Tools.

### Whole mount *in situ* hybridizations

*In situ* hybridizations were performed according to Sive, Grainger, & Harland, 1997 using probes for *Xenopus brachyury (Xbra)* and *Chordin.* Plasmids for probe synthesis were gifts from J. Smith (Smith et al., 1991) and E.M. DeRobertis (Sasai et al., 1994).

### Antibody staining

Cadherin, fibrillin, FN, 12101 or actin stained samples were fixed for 15 min in MBS containing 3.7% formaldehyde and 0.25% glutaraldehyde. For PG and β-catenin staining, samples were fixed in Dent’s solution (80% methanol and 20% DMSO) overnight at 4°C. Dorsal isolates were bleached, stained, and cleared as previously described (Wallingford, 2010). Primary antibodies were used at the following dilutions: PG at 1:100 (*γ*-catenin; BD Biosciences), β-catenin at 1:1000 (C2206; Sigma Aldrich), cadherin at 1:2,000 (XC; polyclonal from Barry Gumbiner), fibrillin at 1:400 (JB3; Developmental Studies Hybridoma Bank [DSHB]), FN at 1:1000 (monoclonal 4H2 and polyclonal 32FJ were developed in the DeSimone lab; 4H2 is available from DSHB), 12101 at 1:10 (DSHB).

### Optical Microscopy

Confocal microscopy was performed on a Nikon C1 microscope, with Nikon Plan Fluor 20x, Plan Apo 60x, and Plan Apo TIRF 100x objectives. Whole embryo images were taken on a Zeiss Stemi SV 6 with an Excelis HDS camera or on a Zeiss SteREO Lumar.V12 with a Zeiss AxioCam MRm.

### Atomic force microscopy

A JPK Instruments NanoWizard 4a AFM head with the CellHesion 200 module mounted on a Zeiss AxioObserver were used for all adhesion measurements. NanoWorld Arrow TL-2 cantilevers were coated with a 20 μg/ml solution of Corning Cell-Tak adhesive according to the manufacturer’s instructions. Cantilevers were calibrated using the thermal noise-based contact-free method built into the JPK software. Dissociated mesendoderm cells were attached to cantilevers maintaining a constant height for 20 seconds using 5 nN of force. After cells were lifted from the dish, they were allowed to adhere for 5 minutes before force measurements were taken. Once the cells were attached to the cantilevers, nascent substrate adhesions were formed by applying 1 nN of force for 5 seconds. Substrates tested were 20 μg/ml of *Bovine* plasma FN, *Xenopus* C-Cadherin-FC, and PLL. Three independent measurements were taken for each cell on a given substrate and statistics were performed on averages of the maximum adhesive strengths for all three measurements.

### Western blots

Embryos were lysed in a modified RIPA buffer and proteins were separated on 6% acrylamide SDS–PAGE gels as described in Bjerke *et al.*, 2014. Western blot membranes were probed using antibodies at the following dilutions: Plakoglobin at 1:1000 (γ-catenin; BD Biosciences), β-catenin at 1:1000 (C2206; Sigma Aldrich), βtubulin at 1:10000 (DM1A; Sigma Aldrich), FN at 1:10000 (4H2; DSHB), C-cadherin at 0.5 μg/ml (6B6; DSHB), integrin β_1_ at 1:1000 (8C8; DSHB).

### Length-to-width ratio and notochord width measurements

Dorsal isolates cut from control and PG morphant embryos were stained for βcatenin to visualize cell outlines. Length-to-width ratios of 5 axial and 5 paraxial mesoderm cells were measured and averaged for each analyzed image. Statistical analyses were performed using these calculated averages. For each image, notochord widths were measured at the anterior and posterior ends as well as in the center. The three width measurements were averaged and used as a single data point for statistical analysis. All measurements were taken from single confocal sections.

### Fibronectin fibrillogenesis measurements

Fibrillogenesis was measured by determining the total area of an immunofluorescence image occupied by fibronectin fibrils. Stage 11 animal caps were stained for FN and imaged using confocal microscopy. Analyses were performed in ImageJ Version 1.50C. Z-stacks were collapsed into z-projections and lower thresholds were set to remove background fluorescence and kept constant across all analyzed images for each experiment. Fluorescent areas corresponding to FN fibrils above threshold levels were measured.

### Injecting embryos with fluorescent FN

*Bovine* plasma fibronectin (Alfa Aesar) was fluorescently labeled with DyLight 550 according to the protocol provided by Thermo Scientific. Embryos were injected with control or PG morpholinos at stage 2. At stage 8, embryos were injected with 20 ng of fluorescent FN directly into the BC. Animal caps were fixed and stained for *Xenopus* FN at stage 11.

### qPCR

RNA was isolated from 10 embryos per condition using the Absolutely RNA Microprep Kit from Agilent. cDNA was synthesized from RNA using the Bioline SensiFAST cDNA Synthesis Kit. cDNA was quantified in the Thermo Scientific PikoReal Real-Time PCR System using the Bioline SensiFAST SYBR No-ROX kit. A plasmid containing the *Xenopus* FN mRNA sequence was used to generate a standard curve for absolute mRNA quantification. Primers used for qPCR reactions were:

*FN* Forward: ACCTCAACTACACAGACTCGAC
*FN* Reverse: AGCTGCCATGGGACAGAATC

### PG rescue construct

A GFP-tagged PG construct (a gift from M.W. Klymkowsky; University of Colorado, Boulder) (Merriam et al., 1997) was mutagenized using site-directed mutagenesis according to the Agilent QuikChange Site-Directed Mutagenesis Kit protocol. A total of 6 silent mutations within the morpholino target sequence were made across 6 amino acids. Mutations were inserted in pairs. PG morpholino target sequence and mutated PG sequence are shown below, with bases targeted for mutagenesis underlined:

PG morpholino target sequence: ATG GAT TT<u>G</u> GG<u>A</u> GA<u>C</u> GT<u>A</u> GT<u>G</u> GA<u>A</u> A
Mutated PG sequence: ATG GAT TT<u>A</u> GG<u>T</u> GA<u>T</u> GT<u>C</u> GT<u>C</u> GA<u>G</u> A

### Surface biotinylation

Surface biotinylation experiment procedures were modified from Gaultier *et al*, 2002. In brief, 25 animal caps per condition were incubated with EZ-Link Sulfo-NHS-LC-Biotin (Thermo) diluted in 0.1x MBS to a concentration of 0.33 mg/ml for 30 minutes at room temperature. Animal caps were transferred to eppendorf tubes and washed five times in 100 mM glycine. Animal caps were then lysed in TBS with 1% Triton X-100, 2 mM phenylmethylsulfonyl fluoride 5 mM EDTA, and Protease Inhibitor Cocktail (Sigma). Yolk and cellular debris was removed by centrifuging lysate for 20 minutes at 14,000 G. Biotinylated proteins were precipitated using NeutrAvidin agarose (Pierce) for one hour at room temperature. NeutravAdin agarose was spun down at 3,000 G for 5 minutes, washed three times with lysis buffer, and biotinylated proteins were removed by boiling for 5 minutes in sample buffer containing SDS.

### Statistics

Graphs were made and statistics were performed using the Prism V5 software package. Wilcoxon match-paired signed rank tests were performed to compare conditions within a given clutch of embryos for western blot analyses. Mann-Whitney tests were used to test for significant differences between two samples across multiple clutches of embryos and experiments, such as embryo and explant lengths or fibronectin fibril formation. Significance was reported according to the following annotation: ns = not significant, *p ≤ 0.05, **p ≤ 0.01, ***p ≤ 0.001.

